# Comparing Lifeact and Phalloidin for Super-resolution Imaging of Actin in Fixed Cells

**DOI:** 10.1101/2020.08.19.248898

**Authors:** Hanieh Mazloom-Farsibaf, Farzin Farzam, Mohamadreza Fazel, Michael J. Wester, Marjolein B. M. Meddens, Keith A. Lidke

## Abstract

Visualizing actin filaments in fixed cells is of great interest for a variety of topics in cell biology such as cell division, cell movement, and cell signaling. We investigated the possibility of replacing phalloidin, the standard reagent for super-resolution imaging of F-actin in fixed cells, with the actin binding peptide ‘lifeact’. We compared the labels for use in single molecule based super-resolution microscopy, where AlexaFluor 647 labeled phalloidin was used in a (d)STORM modality and Atto 655 labeled lifeact was used in a single molecule imaging, reversible binding modality. We found that imaging with lifeact had a comparable resolution in reconstructed images and provided several advantages over phalloidin including lower costs, the ability to image multiple regions of interest on a coverslip without degradation, simplified sequential super-resolution imaging, and more continuous labeling of thin filaments.

## Introduction

Super-resolution methods are optical microscopy methods that surpass the diffraction limit of light, either by manipulating the illumination pattern like STED [1] and SSIM [2] or by stochastic single molecule techniques like (d)STORM [3, 4] and (f)PALM [5, 6]. These methods can achieve resolution that is more than one order of magnitude better than the diffraction limit and provide a tool to investigate biological processes down to the single-molecule level. Equally important in the advancement of super-resolution imaging has been the development of new fluorescent probes. For the stochastic methods, the required intermittent fluorescence (blinking) can be from fluorophores that photo-activate, photo-switch between the on- and off-state or reversibly bind to the target protein. (d)STORM takes advantage of fluorophores that can rapidly switch between on- and off-states for single molecule localization. PAINT uses sparse fluorophore binding to the target structure in order to localize fluorescent events [7]. More recently, DNA-PAINT combined the capabilities of PAINT with programmable DNA hybridization to enable a range of reversible binding mechanisms, leading to higher specificity in target detection as well as higher numbers of individual fluorophores [8, 9].

A common target in super-resolution is the actin cytoskeleton. Actin exists in cells as either free, monomeric globular actin (G-actin) or filamentous actin (F-actin) [10, 11]. The dynamic equilibrium between these states regulates actin filament assembly and disassembly, which in turn controls multiple cellular processes, from cell migration and division to transport of intracellular vesicles. Changes in the regulation of the actin cytoskeleton, whether by mutation of actin itself or actin binding proteins, is associated with a number of diseases [12, 13]. In addition to its biomedical importance, imaging of the actin cytoskeleton is a common benchmark for validation of super-resolution methods. A variety of fluorescent probes for imaging actin have been compared in both live and fixed cells [14, 15], including actin chromobodies [16], utrophin [17], f-tractin [18], silicon rhodamine (SiR)-actin [19], epitope-tagged actin [20], fluorescent protein-actin fusion proteins [21], Affirmers [22], phalloidin as an F-actin-binding reagent [23], and lifeact [24]. In this work, we focus on two actin-labeling reagents, phalloidin and lifeact.

Phalloidin is a heptapeptide derived from the poisonous mushroom *Amanita phalloides* that binds with high specificity to F-actin and prevents filament depolymerization [25, 26]. Direct conjugation of fluorophores to phalloidin makes it a convenient tool for labeling of filamentous actin. However, phalloidin is typically limited to use in fixed cells due to the toxicity induced by its stabilization of the actin filaments and the fact that it is not membrane permeable [23, 27].

Lifeact is a short peptide taken from the first 17 amino acids of the yeast Abp140 and can be expressed in cells linked to a fluorescent protein to visualize F-actin dynamics [24]. It can also be directly synthesized and a milligram amount provides an effectively inexhaustible supply for a typical lab when used for single-molecule imaging. Lifeact has been employed for super-resolution imaging of the actin filaments for a wide range of applications in live cells [28] and in fixed cells [29].

In this article, we compare the quality of lifeact in single molecule super-resolution imaging to phalloidin labeling. To image phalloidin, we used the dSTORM technique [4] where phalloidin was conjugated to AlexaFluor 647 (AF647), a spontaneously photo-switching dye for single-molecule super-resolution imaging [30]. Lifeact was tagged with Atto 655 for a reversible binding approach. We compared lifeact with phalloidin labeling using several metrics including the average resolution of the reconstructed image, apparent filament thickness and continuity, data collection from multiple regions on a coverslip, and for use in sequential super-resolution imaging.

## Methods

### Cell culture and reagents

HeLa cells were cultured in Dulbecco’s Modified Eagle Medium (Life Technologies, Cat No. 10313-v021) supplemented with 10% cosmic calf serum (HyClone), 5U/ml penicillin, 0.05 mg/ml streptomycin, and 2 mM L-glutamine (ThermoFisher, Cat No. 25030081) and maintained at 37 **°**C and 5% carbon dioxide. Rat Basophilic Leukemia (RBL-2H3) cells were cultured in MEM supplemented with 10% heat-inactivated fetal bovine serum, 5 U/ml penicillin, 0.05 mg/ml streptomycin, and 2mM L-glutamine at 37 **°**C and 5% carbon dioxide. For imaging, cells were plated overnight on 25 mm coverslips (1.5, Warner Instruments, #CS-25R15).

### Cell Fixation and Actin Labeling

Samples were fixed using a cytoskeleton-preserving buffer composed of 80 mM Pipes, 5 mM EGTA, and 2 mM MgCl_2_ (PEM) with pH 7.2. Briefly, cells were washed with warm PEM and fixed in 0.6% paraformaldehyde with 0.1% glutaraldehyde and 0.25% Triton diluted in PEM buffer for 60 seconds and followed by a hard fixative buffer including 4% paraformaldehyde and 0.2% glutaraldehyde diluted in PEM for two hours. Cells were washed 2x in PBS, incubated for 10 min in 0.1% NaBH_4_ for 10 min to reduce background fluorescence due to glutaraldehyde, and followed by a 2x wash with PBS. To quench reactive cross-linkers, the samples were incubated in 10 mM Tris for 10 min, followed by 2 washes with PBS. Finally, samples were permeabilized by incubation for 15 min in 5% BSA and 0.05% Triton X-100 diluted in PBS. At the end, samples were washed 1x with PBS and prepared for the labeling process.

To label actin using phalloidin, fixed cells were incubated for an hour in 0.56 μM AF647-conjugated to phalloidin (ThermoFisher, Cat no. A22287) in PBS. The samples were washed once in PBS and placed in dSTORM imaging buffer. The dSTORM buffer included an enzymatic oxygen scavenging system and primary thiol: 50 mM tris, 10 mM NaCl, 10% w/v glucose, 168.8 U/ml glucose oxidase (Sigma, Cat No. G2133), 1404 U/ml catalase (Sigma, Cat No. C9322), and 60 mM 2-aminoethanethiol (MEA) (Sigma-Aldrich, Cat no. M6500-25G) with pH 8.0, to provide a suitable chemical condition for having a photo-switchable AF647 dye.

For lifeact labeling, we used a customized 17-amino-acid peptide conjugated to Atto 655, (sequence: [Atto 655] CMGVADLIKKFESISKEE[COOH] (Bio-Synthesis, Lot No. P5869-1)). For imaging, lifeact was diluted to 0.7 nM in an optimized imaging buffer including 10 mM HEPES, 150 mM NaCl, 10% glucose, and 0.1% BSA with pH 7.0 for labeling the actin filaments.

25 mm coverslips were mounted in an Attofluor cell chamber (Life Technologies, Cat No. A-7816). The corresponding imaging buffer for each approach was added and a clean 25 mm coverslip was used to seal the chamber to avoid oxygen permeation into the buffer.

### Sequential imaging

Sequential super-resolution was performed in a manner similar to that described by Valley *et a*l. [31]. First, actin structures were imaged using either phalloidin-AF647 or lifeact-Atto655. To remove the phalloidin signal, the sample was washed 4x with PBS before exposure to high intensity 638 nm laser light (∼ 4.7 kW/cm^2^) for 5 min to photobleach the sample, followed by a 20 min incubation in 0.1% NaBH_4_ to quench any remaining active AF647 and finally washed 2x with PBS. To remove lifeact-Atto655, the sample was washed 12 x 1 min with 1 mM HEPES, 150 mM NaCl, 5% glucose, and 0.1 % BSA in diH2O. In both cases, the sample was re-labeled with anti-α-tubulin-AF647 at 2.5 μg/ml for an hour in 2% BSA, 0.05% Triton diluted in PBS, followed by 3x wash with 2% BSA and 0.05% Triton in PBS for 5 min each. For data collection, the samples were kept in dSTORM buffer and sealed with a 25 mm coverslip. Re-alignment of the sample was done using a brightfield reference image as described by Valley *et a*l. [31].

### Optical setup

The samples were mounted on the stage of a microscope with a custom-designed chamber holder. The imaging system was built on an inverted microscope (IX71, Olympus America Inc.). A nano positioning stage (Mad City Labs, Nano-LPS100) mounted on an *x-y* manual stage was installed on the microscope for cell location and brightfield registration. A mounted LED with a wavelength of 850 nm (M850L3, Thorlabs) was used for brightfield illumination. Brightfield images were collected on a CMOS camera (Thorlabs, DCC1545M) after reflection by a short-pass dichroic beam splitter (Semrock, FF750-SDi02) and then passing through a single-band bandpass filter (Semrock, FF01-835/70-25). A 638 nm laser (Thorlabs, L638P200) was coupled into a single-mode fiber and reflected by a dichroic beam splitter (Semrock, Di03-R635-t1-25×36) then focused onto the back focal plane of the 1.49 NA objective lens (UAPON 100XOTIRF, Olympus America Inc.). Emission for super-resolution data was collected through a short-pass dichroic beam splitter (Semrock, FF750-SDi02) and a single-band bandpass filter (Semrock, FF624-Di01) on an EMCCD camera (Andor Technologies, iXon DU-860E-CS0-#BV). All the instruments were controlled by custom-written software in MATLAB (MathWorks Inc.) [32]. Imaging was performed with a 638 nm laser at ∼ 4.7 kW/cm^2^ in TIRF (total internal reflection fluorescence) illumination with an approximate exposure time of 40 ms for lifeact and 10 ms for phalloidin (see Exposure time optimization). Brightfield registration was employed to correct for drift after every 3000 frames for phalloidin experiments and 2000 frames for lifeact experiments, as described in [33].

### Data analysis

All analyses were performed using a custom-written single molecule analysis (SMA) software package in MATLAB combined with the Statistics Toolbox (The MathWorks, Inc) and DIPimage [34]. For 2D raw data, single emitters in each frame were identified as single localized spots. The parameters of each emitter, which includes positions, total photon count, background photon counts and their standard errors were computed by maximum likelihood estimation on a GPU [35]. After applying a threshold on maximum background photon counts of 200, minimum photon counts per frame per emitter of 250, and a data-model hypothesis test [36] with a cutoff p-value at 0.01, a well-defined set of localizations was reconstructed to generate a super-resolution image. To eliminate outliers, a nearest neighbor filter was employed to remove those localizations having less than 4 neighbors within 15 nm (see S1 Fig).

### Fourier Ring Correlation (FRC) analysis

To estimate the resolution from the super-resolution images of phalloidin and lifeact, we used Fourier Ring Correlation (FRC) [37], which measures the average resolution over a single super-resolution image. In this method, the localized single emitters of a super-resolution image are divided into two statistically independent subsets. The image resolution is defined as the inverse of the spatial frequency when the FRC curve drops below a threshold of 1/7. Spurious correlations were removed by estimating the number of times an emitter was localized on average (*Q*) assuming Poisson statistics. All analyses were accomplished using the MATLAB software developed by Nieuwenhuizen *et a*l. [37] found at http://www.diplib.org/add-ons.

### Exposure time optimization

To acquire optimized super-resolution data, the exposure times were adjusted to be the same as the average blinking on-time. The on-time periods have an exponential distribution with an average of 1/k_off_, where k_off_ is the rate of emitters going from on to off in units of 1/ms (S2 Fig). For the lifeact approach, data sets were collected at various exposure times. To estimate the average on-time of blinking events, the localizations from the same blinking events were connected across consecutive frames and the on-time histogram was fit to an exponential distribution using a Poisson noise model. The likelihood is as follows

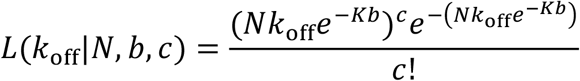

where *b, c* and *N*, respectively, stand for the number of localizations per binding event, the frequency of blinking events with a specific number of connected localizations, and the total number of localizations. The average on-time of the blinking events is then given by τ=Δ*T*/*k*_off_, where Δ*T* is the exposure time for data acquisition.

### Thickness and continuity assessment of actin filaments in reconstructed images

To evaluate the thickness and continuity of actin filaments in reconstructed images, regions of interest (ROIs) containing a segment of filament were selected manually. The selected segments were rotated to be along the *x*-axis to facilitate subsequent comparisons. The aligned actin segments were used to find the sum of normalized intensity distributions along and perpendicular to the *x-*axis to assess the continuity and thickness of the actin filaments, respectively. The resulting intensity distributions perpendicular to the *x*-axis shows the cross section of actin filaments, and their widths at half maximum were reported as actin thicknesses.

## Results

### Resolution in super-resolution images

We employed the two actin labeling approaches using phalloidin and lifeact and compared the super-resolution images of actin structures in fixed HeLa and RBL-2H3 cells. To evaluate the resolution in each treatment, we used Q-corrected Fourier Ring Correlation (FRC), which measures the average resolution over a single super-resolution image [37]. The Q-correction removes the effect of repeat blinking events that can artificially improve the resolution measure. The reversible binding of lifeact generates a small percentage of spurious localizations from transient non-specific binding. These appear as isolated localizations away from any filaments. Localizations that did not have at least four other localizations within 15 nm were removed before FRC analysis (see Methods, S1 Fig).

Actin filaments of HeLa cells are shown in Fig 1A and 1B. The average resolution measurements for the lifeact and phalloidin images were 41.10 +-0.73 nm, and 41.37 +-0.4 nm, respectively. We repeated the same scenario for RBL-2H3 cells (Fig 1C, 1D) and obtained average resolutions of 43.73 +-0.57 nm, and 35.86 +-0.37 nm for lifeact and phalloidin, respectively. The FRC curve estimates the average resolution of a super-resolution image as the inverse of the spatial frequency when the curve crosses the 1/7 line (Fig. 2, red line, see also Methods). As depicted in Fig. 2A (2B) for three HeLa (RBL-2H3) cells from different samples, phalloidin produced an equal or slightly improved average resolution compared to the lifeact.

**Fig 1.**
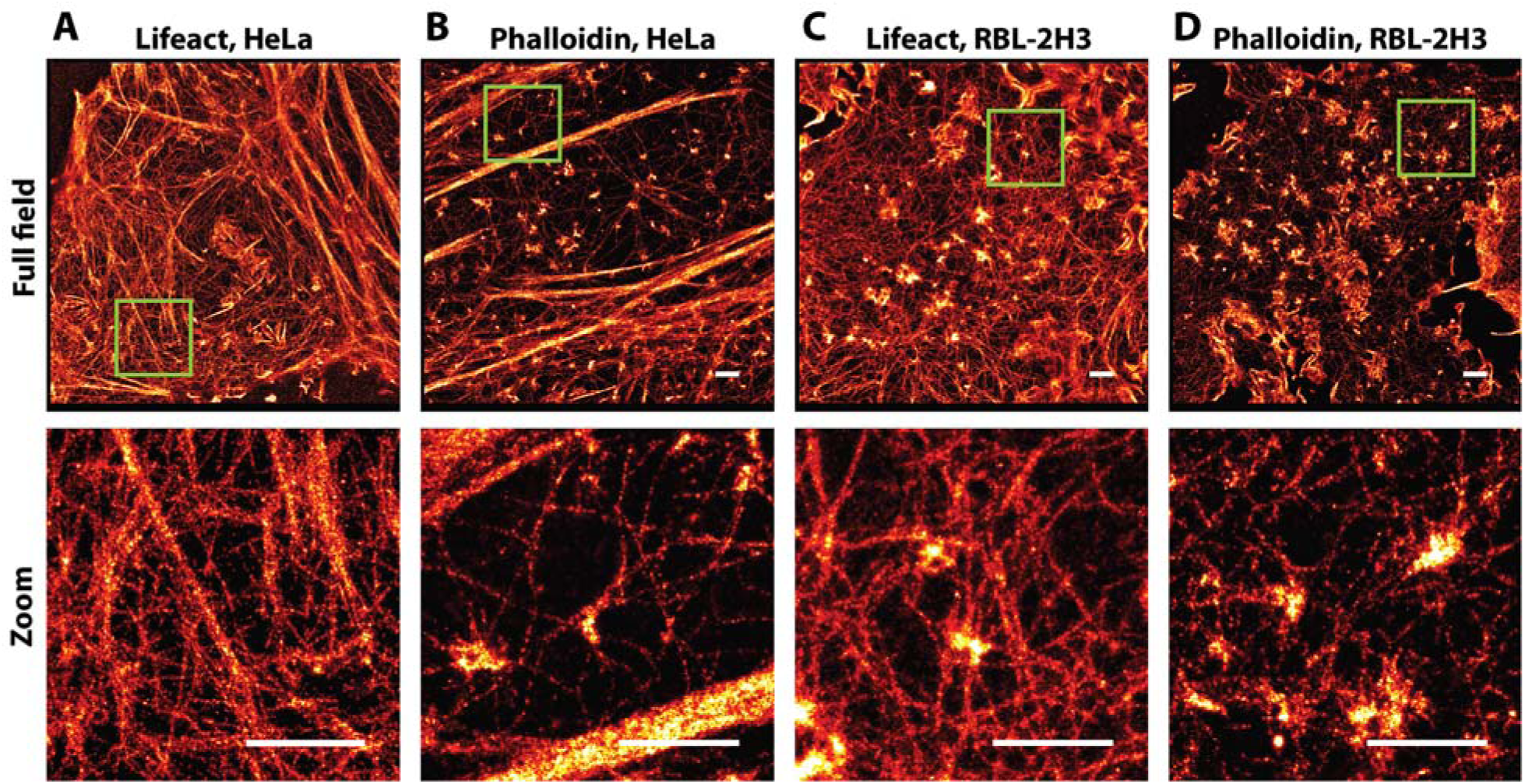
Super-resolution images of F-actin. (A) F-actin labeled with lifeact-Atto655 in a Hela cell image (top), zoomed-in image (bottom). (B) F-actin labeled with phalloidin-AF647 in a Hela cell (top), zoomed-in image (bottom). (C) F-actin labeled with lifeact-Atto655 in a RBL-2H3 cell image (top), zoomed-in image (bottom). (D) F-actin labeled with phalloidin-AF647 in a RBL-2H3 cell image (top), zoomed-in image (bottom). Scale bars, 1 μm.

**Fig 2.**
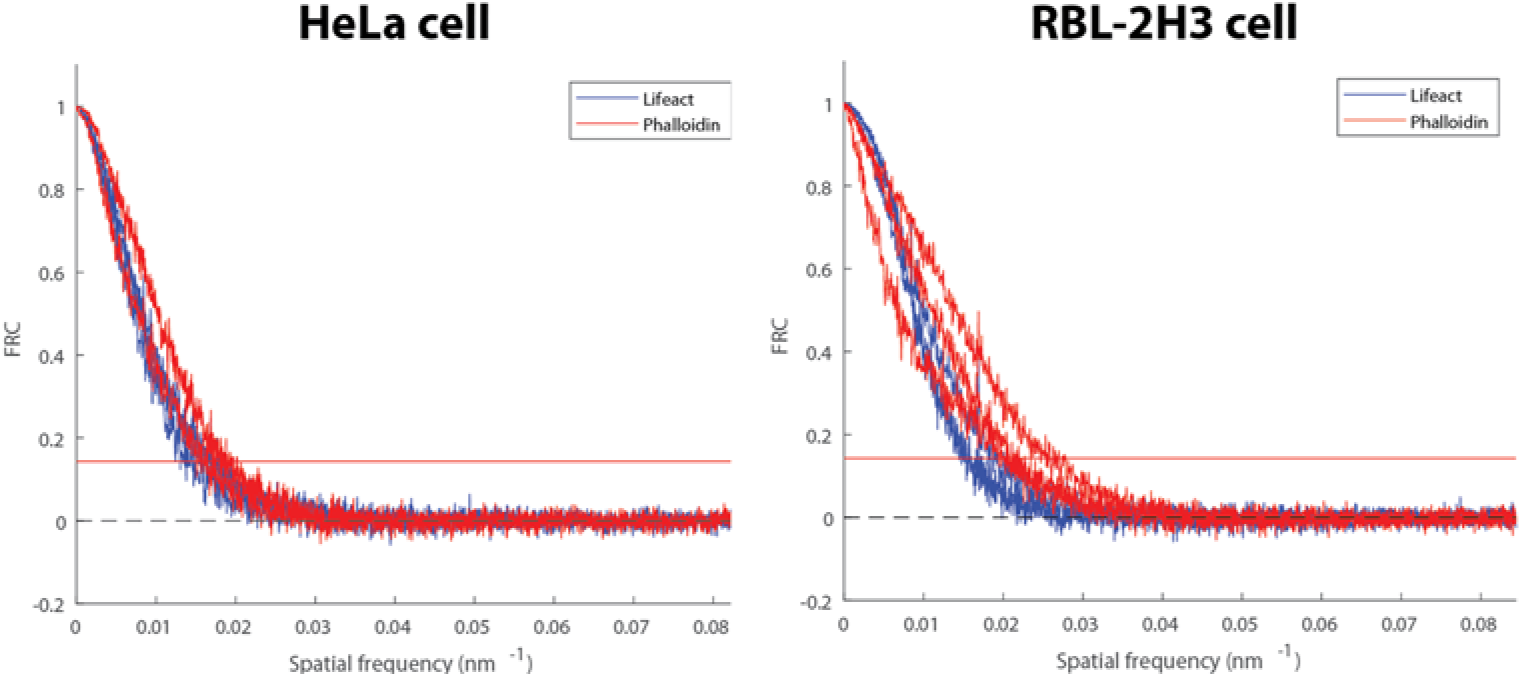
FRC curves for super-resolution images of F-actin. (Left) FRC curve of F-actin super-resolution images in HeLa cells from three samples of independent experiments for lifeact (blue) and phalloidin (red). (Right) FRC curves of F-actin super-resolution images in RBL-2H3 cells from three samples of independent experiments for lifeact (blue) and phalloidin (red). The horizontal red line at FRC = 1/7 indicates the value of the average resolution, which is defined as the inverse of the spatial frequency at this point.

### Filament thickness and continuity

We estimated the thickness and continuity of actin filaments found in images from each condition. As illustrated in Fig. 3 with small green boxes, several thin filaments were chosen from the reconstructed super-resolution image for lifeact (Fig 3A) and phalloidin (Fig 3B). The reconstructed thickness was found to be approximately 24 nm at FWHM (Full Width Half Maximum) for both methods (see Methods, Fig 3C). To evaluate the continuity of observed actin filaments for each strategy, the normalized intensity along thin filaments is shown in Fig 3D. The individual selected filaments for lifeact show continuity of localized single molecules along the filament length (Fig 3D, blue graph). In contrast, for the phalloidin experiment, the normalized intensity distribution along the actin filaments falls to zero more frequently, signifying more discontinuous labeling (Fig 3D, red graph). The location of bound phalloidin is determined at the sample preparation stage and therefore the resolution and continuity of thin filaments are limited by the initial labeling. Lifeact can continuously and stochastically sample actin filaments and therefore longer imaging times can be used to more completely label structures (S3 Fig).

**Fig 3:**
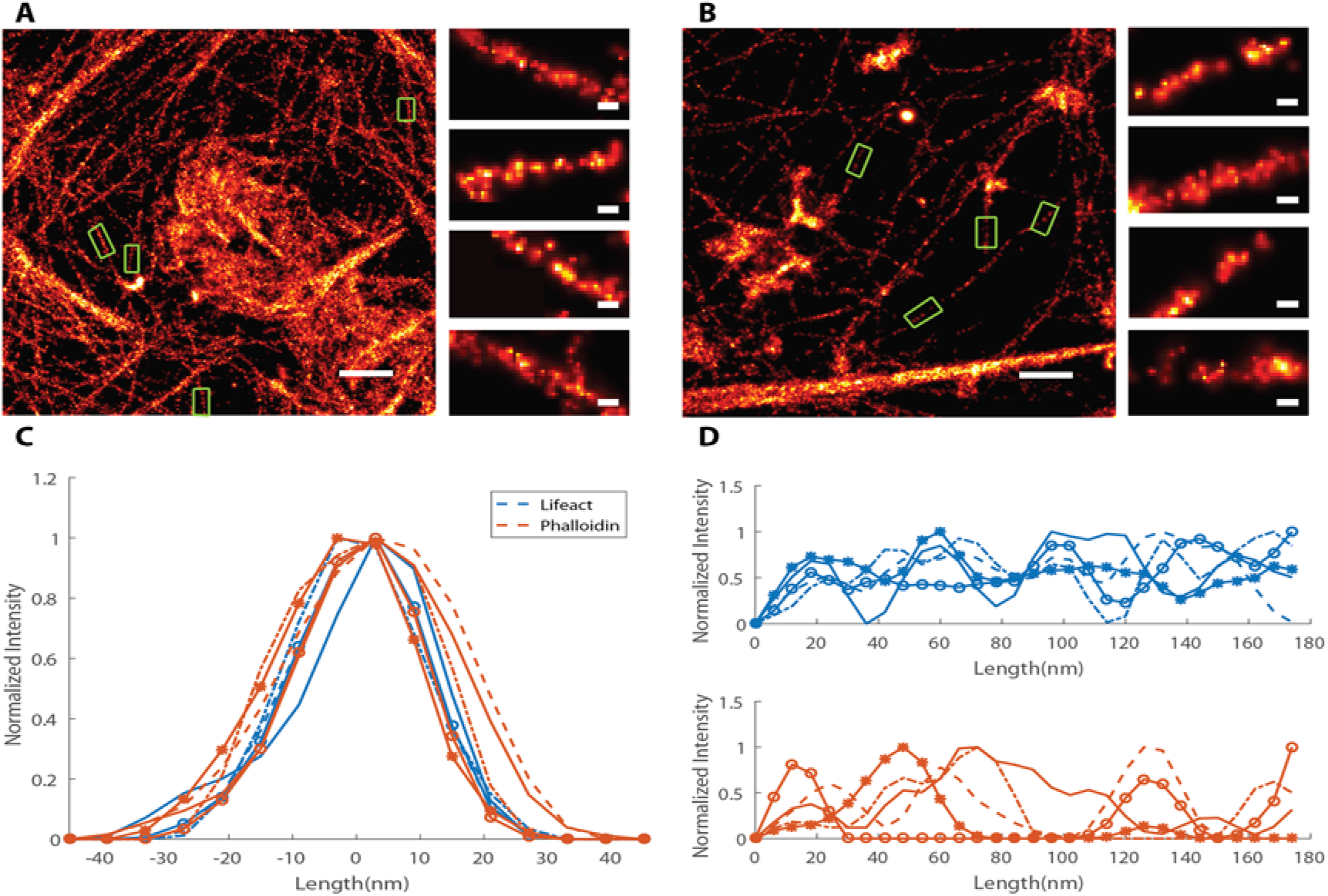
Comparison of lifeact and phalloidin for individual actin filaments. (A) Super-resolution image from lifeact tagged sample. (B) Super-resolution image from phalloidin tagged sample. The green boxes show several selected filaments. (C) The normalized intensities of the selected filaments through the filament cross-section. (D) The normalized intensities along the selected filaments, for the lifeact (top) and phalloidin (bottom) methods. The scale bars in full field images are 500 nm. The scale bars in zoomed regions are 30 nm.

### Multiple dataset collection

It is often desired to collect data from multiple cells or regions on one coverslip. To investigate this aspect, we imaged actin structures for an extended period using both labeling methods. We found that the lifeact approach provided a near constant number of localizations per frame, whereas the number of localizations per frame with phalloidin imaging dropped considerably even during the imaging of the first cell on a coverslip (S4 Fig). This can be explained by reversible binding of lifeact from a large pool of lifeact in the buffer whereas phalloidin suffers from both photobleaching and dissociation over the timescale of hours. Fig 4 depicts the reconstructed super-resolution image for the lifeact approach where imaging began after 10 min (Fig 4A) or 17 hr (Fig 4B) of adding the imaging buffer. We also collected data with the phalloidin approach with imaging starting after 5 min (Fig 4C) or 2 hr (Fig 4D). Lifeact imaging showed no decrease in performance for the later data collection while phalloidin suffered from reduced phalloidin labeling due to dissociation before the start of imaging.

**Fig 4.**
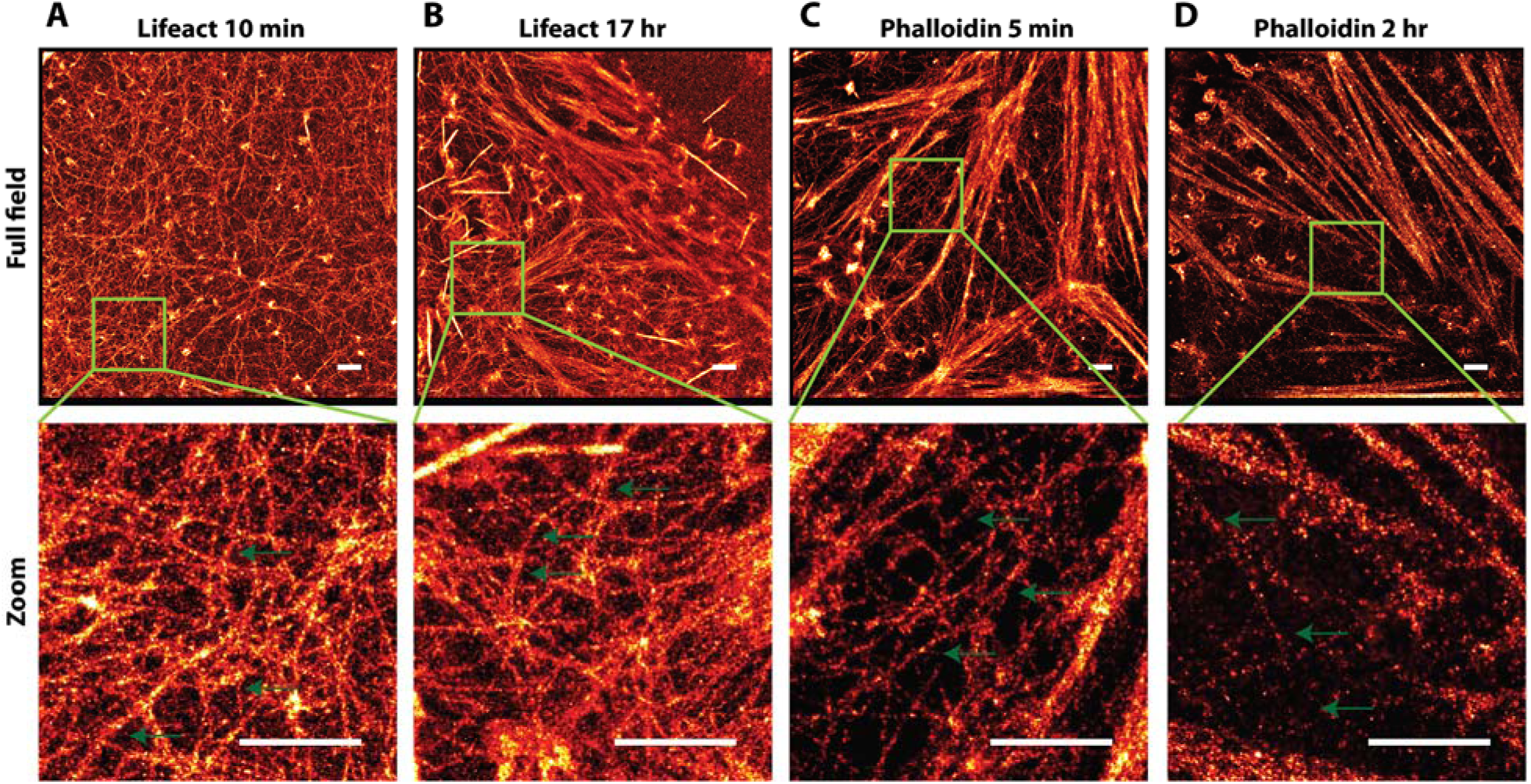
Super-resolution of actin filaments in HeLa cells at various time-points after the start of imaging. (A) Super-resolution of F-actin labeled with lifeact-Atto655 in the full field image (top) and the zoomed-in image (bottom); data collection was started at 10 min after adding the imaging buffer;. (B) Super-resolution of F-actin labeled with lifeact-Atto655 in the full field image (top) and the zoomed-in image (bottom); data collection was started at 17 hr after adding the imaging buffer. (C) Super-resolution of F-actin labeled with phalloidin-AF647 in the full field image (top) and the zoomed-in image (bottom); data was collected after adding the imaging buffer within 5 min. (D) Super-resolution of F-actin labeled with phalloidin-AF647 in the full field image (top) and the zoomed-in image (bottom); data was collected after adding the imaging buffer within 2 hr. The green arrows point to some thin filaments to show their continuities. The scale bar is 1 μm.

### Sequential imaging

Imaging of multiple structures within the same cell is of interest in order to reveal a spatial correlation between the cellular structures. Here, we compared phalloidin and lifeact strategies in their capacity for sequential super-resolution imaging [31]. As depicted in Fig 5A, the first round of labeling was to detect actin using lifeact or phalloidin. After removal of the actin-binding probe (see Methods), microtubules were imaged with dSTORM in the same samples (Fig 5B). The green boxes in Fig 5A showcase bright actin bundles, where abundance of actin-binding markers were detected before being removed completely for the second round of imaging on microtubules (Fig 5B). Fig 5C shows the overlay of actin filaments and microtubules in the subregions indicated in Fig 5B.

**Fig 5.**
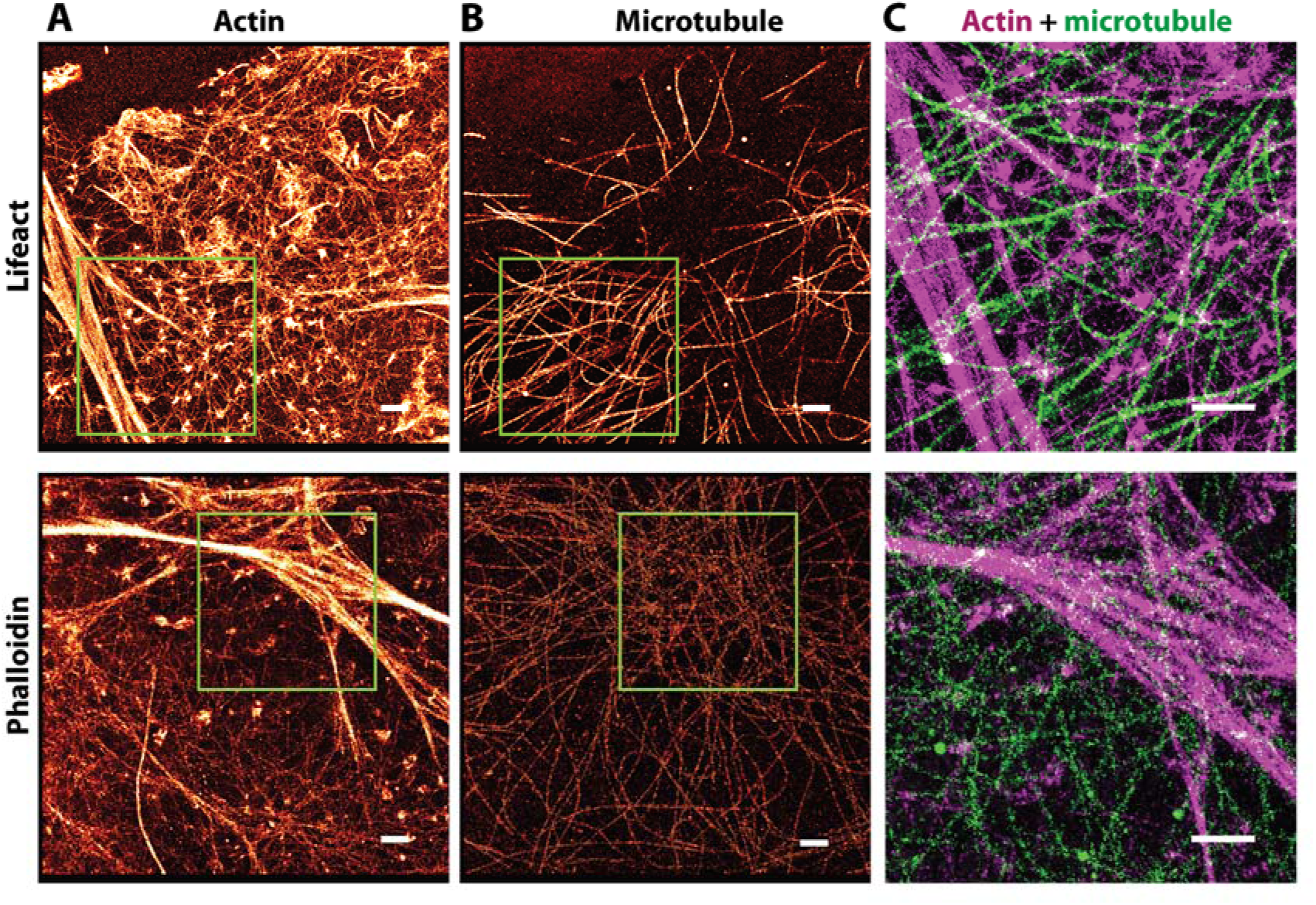
Sequential imaging of actin structures and microtubules in HeLa cells. (A) Super-resolution image of F-actin labeled with lifeact-Atto655 (top) and phalloidin-AF647 (bottom). (B) Super-resolution image of microtubules of the same cell. (C) Overlay of actin structures (red) and microtubules (green) from the subregion selected by the green box when lifeact (top) or phalloidin (bottom) were used for actin imaging. The scale bar is 1 μm.

## Discussion

In evaluating super-resolution imaging results of the lifeact and phalloidin labeling methods, we considered four aspects: image resolution, filament thickness and continuity, multiple dataset collection, and capability for sequential imaging. The resolution as measured by FRC analysis showed that the average resolution in the case of lifeact is comparable or to that with phalloidin, with phalloidin having a small advantage in the RBL cells but not in the HeLa cells. It is important to note that the FRC resolution measure gives an average resolution over the image, but cannot directly assess the quality of labeling.

To evaluate labeling quality, we quantified individual filaments in the reconstructed super-resolution images by looking at filament thickness and continuity. Single filament analysis revealed the average thickness is 24 nm for both labeling approaches, but suggested the continuity of actin filaments improves in the lifeact method. For actin bundles where the number of binding sites are much higher than for single filaments, the reconstructed super-resolution images appear as bright regions in both methods; however, for single filaments, the lifeact method results in a more continuous visualization of the actin filaments. It has been shown that targeting the actin binding sites depends on the biochemical structure of each actin-detecting probe and the various actin architectures [15]. For example, lifeact fails to detect some fine structures such as filopodia in mesenchymal cells [38] or cofilin-bound actin in STHdh cells [39] whereas phalloidin does not bind to filaments with less than seven actin subunits [40].

The blinking mechanism in the lifeact method depends on the binding/unbinding of the peptide, and benefits from a photostable dye, such as Atto655. This provides for consistent localizations over long imaging periods as the probe is replenished from the buffer pool. However, the background fluorescence from the fluorescent, diffusing lifeact requires a TIRF or near-TIRF modality to reduce background, restricting imaging to near the coverslip. For phalloidin labeling, the blinking mechanism is provided by light-activated photoswitching of the AF647 label, which undergoes photobleaching during data collection [31]. With lifeact, super-resolution images collected several hours after adding the imaging buffer showed no degradation of labeling, enabling multiple dataset collection from one coverslip. With phalloidin, the best image quality was always achieved from the first region imaged, with imaging performed just after washing out the free label. Subsequent regions suffered from reduced labeling from phalloidin dissociation.

Both methods were successful with the use of sequential labeling to image actin and microtubules within the same cell. Lifeact had the advantage that the probe could simply be washed out before microtubule labeling. After imaging phalloidin, the additional steps of photobleaching and quenching were required to eliminate crosstalk with the first imaging cycle.

In summary, we found that lifeact provides a reliable and inexpensive alternative to phalloidin for actin visualization in fixed cells, particularly for super-resolution imaging near the coverslip. Imaging with lifeact additionally confers several advantages over phalloidin including more continuous labeling of filaments, the ability to image multiple cells per sample and simplification of sequential super-resolution imaging.

## Acknowledgments

This work was supported by NIH grants R01GM109888, 1R21EB019589 and the New Mexico Spatiotemporal Modeling Center (NIH P50GM085273). We gratefully acknowledge the use of the University of New Mexico Comprehensive Cancer Center fluorescence microscopy core, as well as the NIH P30CA118100 support for these cores. We thank Shayna Lucero for assistance with cell culture and Diane Lidke for suggestions on the manuscript.

---

**S1 Fig.**
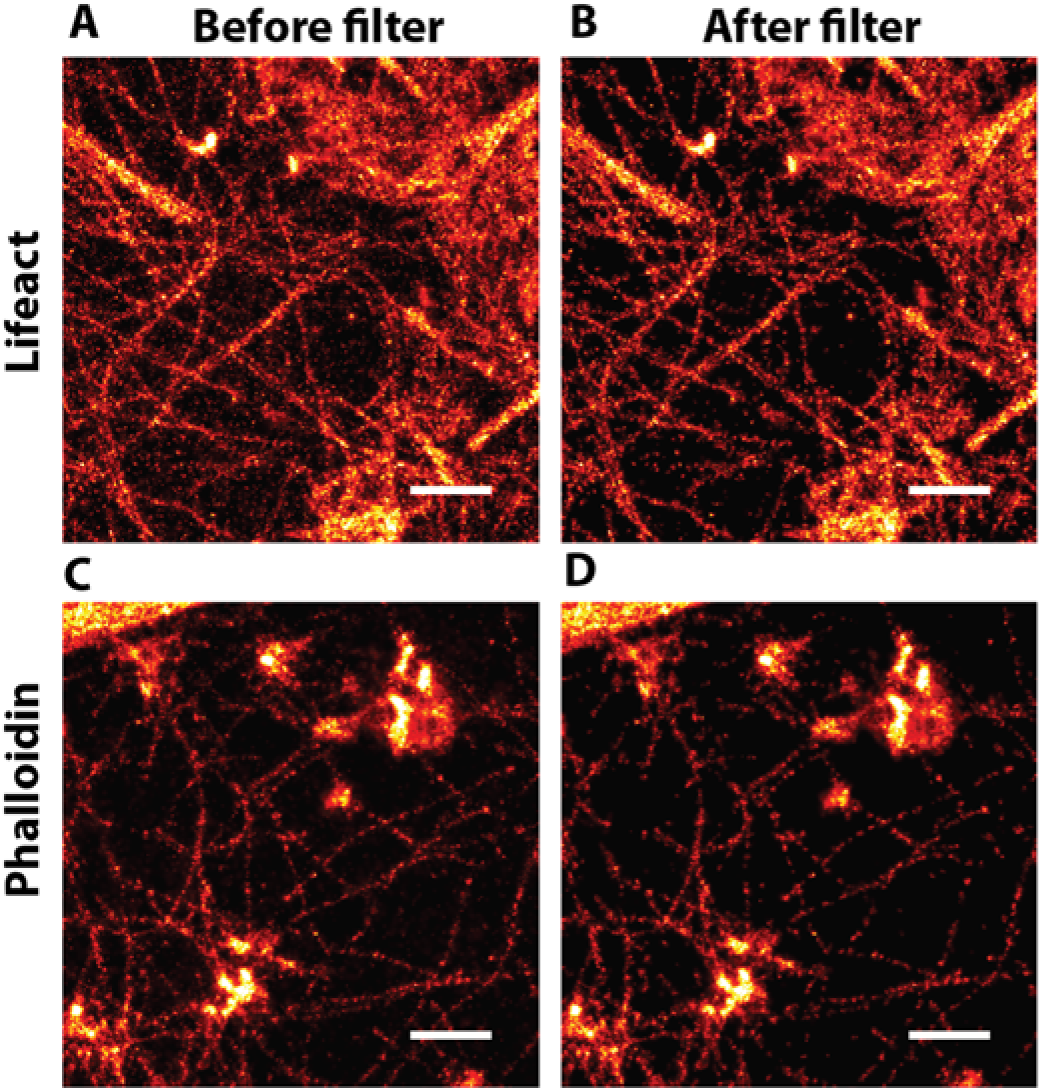
Nearest neighbor filtering to remove nonspecific localizations. (A) Reconstructed super-resolution images of actin filaments in HeLa cells labeled with lifeact-Atto655 (top) and phalloidin-AF647 (bottom). (B) Super-resolution images of actin filaments in HeLa cells labeled with lifeact-Atto655 (top) and phalloidin-AF647 (bottom) after removing localizations that had less than 4 other localizations within 15 nm. Scale bars are 500 nm.

**S2 Fig.**
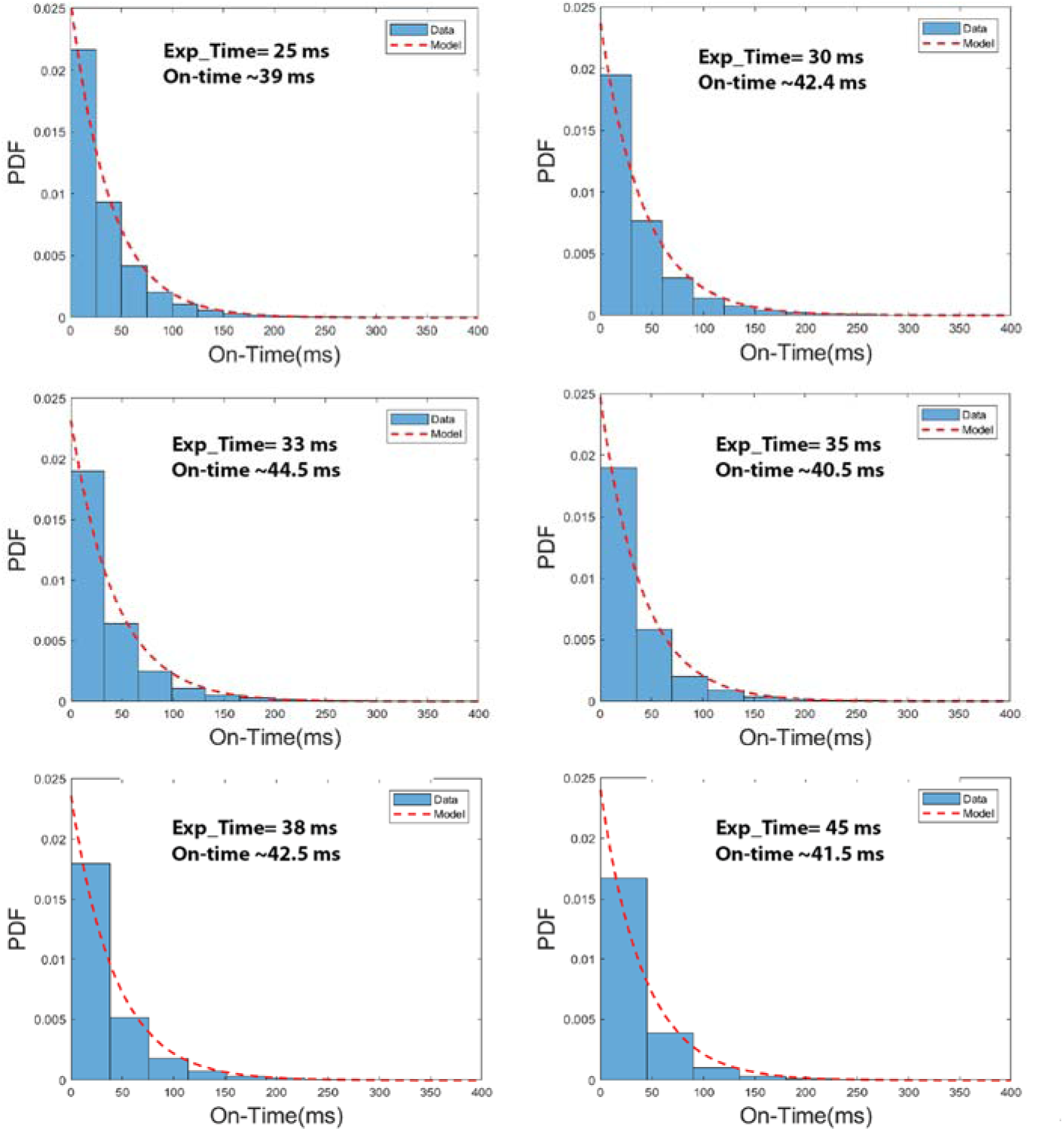
Estimation of average binding time for lifeact-Atto655. Data was collected with six exposure times: 25 ms, 30 ms, 33 ms, 35 ms, 38 ms, and 45 ms. The distribution of the number of connected localizations per binding event (blue) were fit with an exponential model (red dashed line) to extract a binding lifetime.

**S3 Fig.**
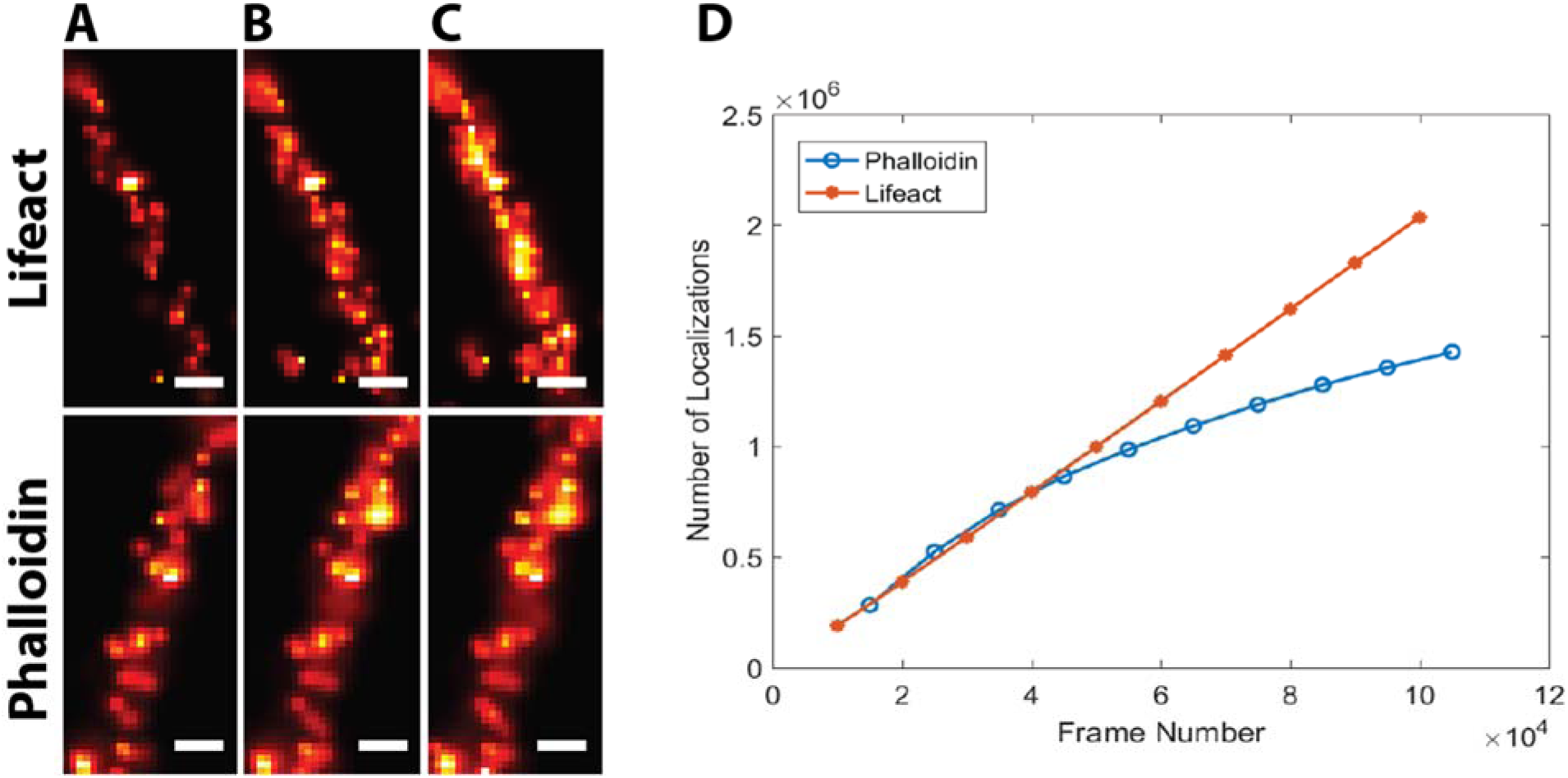
Cumulative number of localizations in actin filaments. Single filaments were cropped from the reconstructed super-resolution images from lifeact-Atto655 (top) and phalloidin-AF647 (bottom). The filaments were reconstructed using localizations from the first (A) 30,000, (B) 60,000 and (C) 100,000 frames. The scale bars are 20 nm. (D) The cumulative number of localizations per frame from data collected using lifeact (red) and phalloidin (blue). Data is from HeLa cells.

**S4 Fig.**
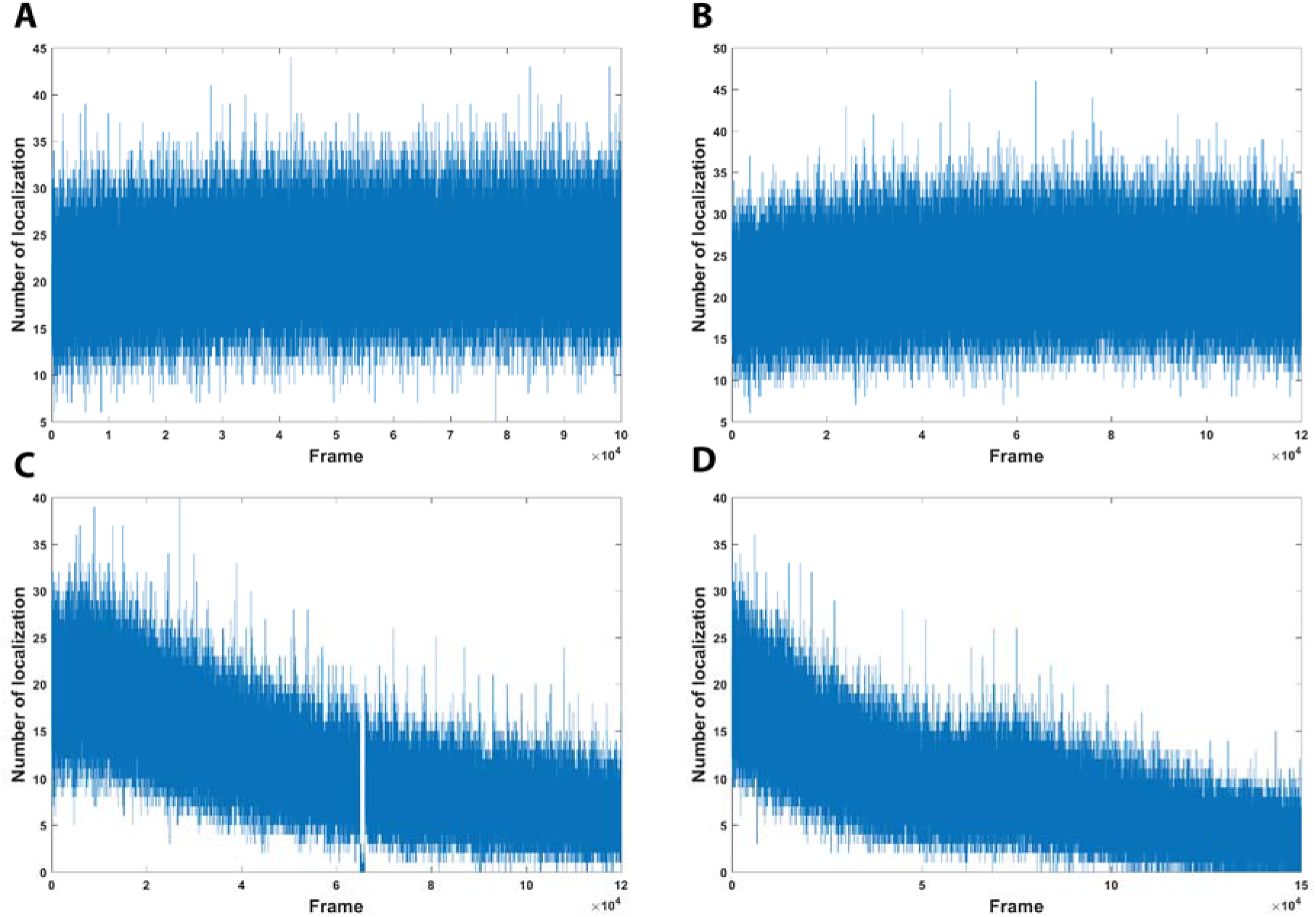
Localizations per frame. Plots show the number of localized single molecules per frame for complete data collections for individual cells. (A) HeLa cells labeled with lifeact-Atto655. (B) RBL-2H3 cells labeled with lifeact-Atto655. (C) HeLa cells labeled with phalloidin-AF647. (D) RBL-2H3 cells labeled with phalloidin-AF647.

